# Laminar-specific interhemispheric connectivity mapping with bilateral line-scanning fMRI

**DOI:** 10.1101/2021.03.08.433876

**Authors:** Sangcheon Choi, Yi Chen, Hang Zeng, Bharat Biswal, Xin Yu

**Affiliations:** Max Planck Institute for Biological Cybernetics, Tübingen, Baden-Württemberg, Germany; Graduate Training Centre of Neuroscience, University of Tübingen, Tübingen, Baden-Württemberg, Germany; Department of Biomedical Engineering, NJIT, Newark, NJ, USA; MGH/MIT/HMS Athinoula A. Martinos Center for Biomedical Imaging, Department of Radiology, Harvard Medical School, Massachusetts General Hospital, Charlestown, Massachusetts, USA

**Keywords:** interhemispheric functional connectivity, callosal projection, global neuromodulation, high-resolution fMRI

## Abstract

Despite extensive studies detecting blood-oxygen-level-dependent (BOLD) fMRI signals across two hemispheres to present cognitive processes in normal and diseased brains, the role of corpus callosum (CC) to mediate interhemispheric functional connectivity remains controversial. Several studies show maintaining low-frequency fluctuation of resting-state (rs)-fMRI signals in homotopic brain areas of acallosal humans and post-callosotomy animals, raising the question: how can we specify the circuit-specific rs-fMRI signal fluctuation from other sources? To address this question, we have developed a bilateral line-scanning fMRI (BiLS) method to detect bilateral laminar BOLD fMRI signals from symmetric cortical regions with high spatial (100 μm) and temporal (100 ms) resolution in rodents under anesthesia. In addition to ultra-slow oscillation (0.01-0.02 Hz) patterns across all cortical layers, a layer-specific bilateral coherence pattern was observed with a peak at Layer (L)2/3, where callosal projection neurons are primarily located and reciprocal transcallosal projections are received. In particular, the L2/3-specific coherence pattern showed a peak at 0.05 Hz based on the stimulation paradigm, depending on the interhemispheric CC activation. Meanwhile, the L2/3-specific rs-fMRI coherence was peaked at 0.08-0.1Hz which was independent of the varied ultra-slow oscillation patterns (0.01-0.02 Hz) presumably involved with global neuromodulation. This work provides a unique laminar fMRI mapping scheme to characterize the CC-mediated evoked fMRI and frequency-dependent rs-fMRI responses, presenting crucial evidence to distinguish the circuit-specific fMRI signal fluctuations across two hemispheres.

**Significance statement:** Laminar fMRI is a promising method to better understand neuronal circuit contribution to functional connectivity (FC) across cortical layers. Here, we developed a bilateral line-scanning fMRI method, allowing the detection of laminar-specific BOLD-fMRI signals from homologous cortical regions in rodents with high spatial and temporal resolution. Laminar coherence patterns of both evoked and rs-fMRI signals revealed that CC-dependent interhemispheric FC is significantly strong at Layer 2/3, where callosal projection neurons are primarily located. The Layer 2/3-specific rs-fMRI coherence is independent of ultra-slow oscillation based on global neuromodulation, distinguishing the circuit-specific rs-fMRI signal fluctuation from different regulatory sources.

## INTRODUCTION

Resting-state (rs-) fMRI, as a non-invasive neuroimaging method, detects functional connectivity (FC) in the brain by measuring fMRI signal fluctuations during rest [1–5].The fMRI signal oscillates at a low-frequency range less than 0.1 Hz, representing strong correlations among functional modules, *e.g.,* symmetric cortices of two hemispheres [6, 7]. Corpus callosum (CC), as major fibers connecting homologous cortical areas of two hemispheres, is considered to mediate bilateral FC [7, 8]. Although significantly diminished bilateral FC has been reported in acallosal human brains [9–12] and post-callosotomy rodents [13–15], several reports have demonstrated nearly intact interhemispheric FC in individuals with callosotomy [16–19]. Also, a bilateral EEG study of humans [20] and a rs-fMRI study of monkeys [21] further demonstrated that after complete callosotomy the bilateral FC is reduced but exist, attributing to the brain network coordination via subcortical mechanism. Consequently, the controversial role of CC underlying bilateral FC raises a fundamental question: how does brain structural wiring contribute to resting-state FC? In recent studies, emerging evidence has shown that subcortical neuromodulatory projections mediate brain state changes, contributing to global and region-specific fMRI signal fluctuation [22–30]. It is thus plausible that the bilateral FC can be regulated by both callosal connections and subcortical neuromodulatory projections.

High field fMRI reveals laminar-specific responses to either bottom-up or top-down tasks, indicating neuronal circuit-based laminar specificity in human [31–35] and animal brains [36–40]. In contrast, the laminar-specific fMRI signal correlation/coherence patterns, which can illustrate underlying neuronal circuits for FC, have not been thoroughly investigated using conventional fMRI methods given the limited spatiotemporal resolution. Yu et al. have developed a line-scanning fMRI method to substantially improve spatial (50 μm) and temporal (50 ms) resolution with a line profile across different cortical layers in rat brains [38]. The ultra-fast sampling scheme of this method avoids the aliasing of the cardiorespiratory cycles over low-frequency fluctuation of rs-fMRI signals. Whereas, most slow sampling methods need retrospective correction for human brain mapping [41–47] and aliasing effect remains a challenging issue for rodent fMRI [48–52]. Meanwhile, the ultra-high spatial resolution provides a unique advantage to map distinct laminar BOLD signals with peripheral or optogenetic stimulation across the 1-2 mm rodent cortex [38, 39]. Given the thickness of gray matter of human brains is in the comparable scale of the rodent cortex ranging from 1-4 mm [53], the line-scanning fMRI enables the detailed layer-specific analysis of fMRI signals. Recently, both line-scanning BOLD and diffusion cortical mapping has been implemented to investigate layer-specific anatomical and evoked hemodynamic features in human brains [54–57]. Also, a recent animal fMRI study co-registered the brainwide rs-fMRI correlation pattern and 3D Allen mouse brain atlas, presenting axonal tracing-base layer specific neuronal connections underlying the default mode network [58]. To date, however, no direct measurements of laminar fMRI signals have been performed on symmetric cortices of two hemispheres to differentiate circuit-specific regulatory sources with high spatiotemporal resolution.

To study laminar-specific intrahemispheric FC, we developed a multi-slice line-scanning fMRI method to detect evoked and rs-fMRI signals from multiple line profiles with ultra-high spatiotemporal resolution [59]. Here, we applied a bilateral line-scanning (BiLS) method to record laminar fMRI signals in bilateral forepaw somatosensory cortex (FP-S1) of anesthetized rats. In both evoked and resting states, we detected ultra-slow oscillation (0.01-0.02 Hz) across all cortical layers, which can be directly differentiated from the callosal circuit-specific bilateral coherent oscillation at Layer (L) 2/3, where callosal projection neurons are mainly located [60, 61]. For experiments with stimulation (20 s per epoch with 4 s on and 16 s off), the coherent frequency of bilateral BOLD signals was detected at 0.05 Hz, but during rest, the coherent frequency of bilateral rs-fMRI signal fluctuation was detected at 0.08-0.1 Hz. Our findings demonstrate that our BiLS method enables the decomposition of frequency-specific interhemispheric low-frequency fluctuations, revealing two independent laminar-specific coherent bandwidths *(i.e.,* 0.01-0.02 Hz versus 0.05 Hz for evoked fMRI or 0.08-0.1 Hz for rs-fMRI) driven by different neuronal sources.

## RESULTS

### Mapping the evoked BOLD fMRI signals with bilateral line-scanning fMRI

We first extended the line-scanning fMRI method from single slice-based unilateral to bilateral BOLD-fMRI mapping with BiLS on the forepaw somatosensory cortex (FP-S1).**Fig. 1A** depicted the line-scanning scheme with two saturation slices to control the width of a field-of-view (FOV) and to suppress signals outside the FOV. **Fig. 1B** demonstrated FP-S1 BOLD responses across different cortical layers from the representative trial and averaged 2D fMRI maps. **Fig. 1C** demonstrated the bilateral FP-S1 BOLD signals upon left forepaw electrical stimulation with BiLS method, showing dynamic BOLD responses as a function of time in the FP-S1 of both hemispheres. In contrast to highly robust BOLD signals in right FP-S1 (4 s on/16 s off for each 20 s epoch), the ultra-slow oscillation of baseline fMRI signals was observed in the left FP-S1 ipsilateral to stimulation (**Fig 1C and D**). Power spectral density (PSD) analysis of both sides showed the 0.05 Hz peak (**Fig. 1F**, magenta arrows) given the 20 s stimulation paradigm for each epoch, and PSD peaks of the ultra-slow fluctuation (<0.02 Hz) were detected in both FP-S1 (**Fig. 1F**), which has been previously reported in anesthetized rats during rest [62]. This result demonstrated the feasibility of BiLS to acquire the interhemispheric laminar-specific fMRI signals with high spatiotemporal resolution. It should also be noted that the fast-sampling of the line-scanning scheme enables estimating the impact of cardiac and respiratory cycles on the rs-fMRI signal fluctuation. We had monitored respirations and blood pressures during fMRI experiments to calculate the spectrogram of the cardiorespiratory responses presenting 5-8 Hz cardiac and 1 Hz respiratory cycles of the anesthetized rats under ventilation (**Fig. S1 A and B**). Meanwhile, the potential cardiac-related pulsation effect could be identified at the superficial L1, but diminished in the deeper layers (**Fig. S1 C**) [63, 64]. Also, due to the usage of muscle relaxer during ventilation [48], the artifacts due to respiration-related B0 fluctuations [65, 66] was negligible in the line-scanning fMRI signals. This result further supported the elucidation of circuit-specific fMRI signals with BiLS method.

**Figure 1.**
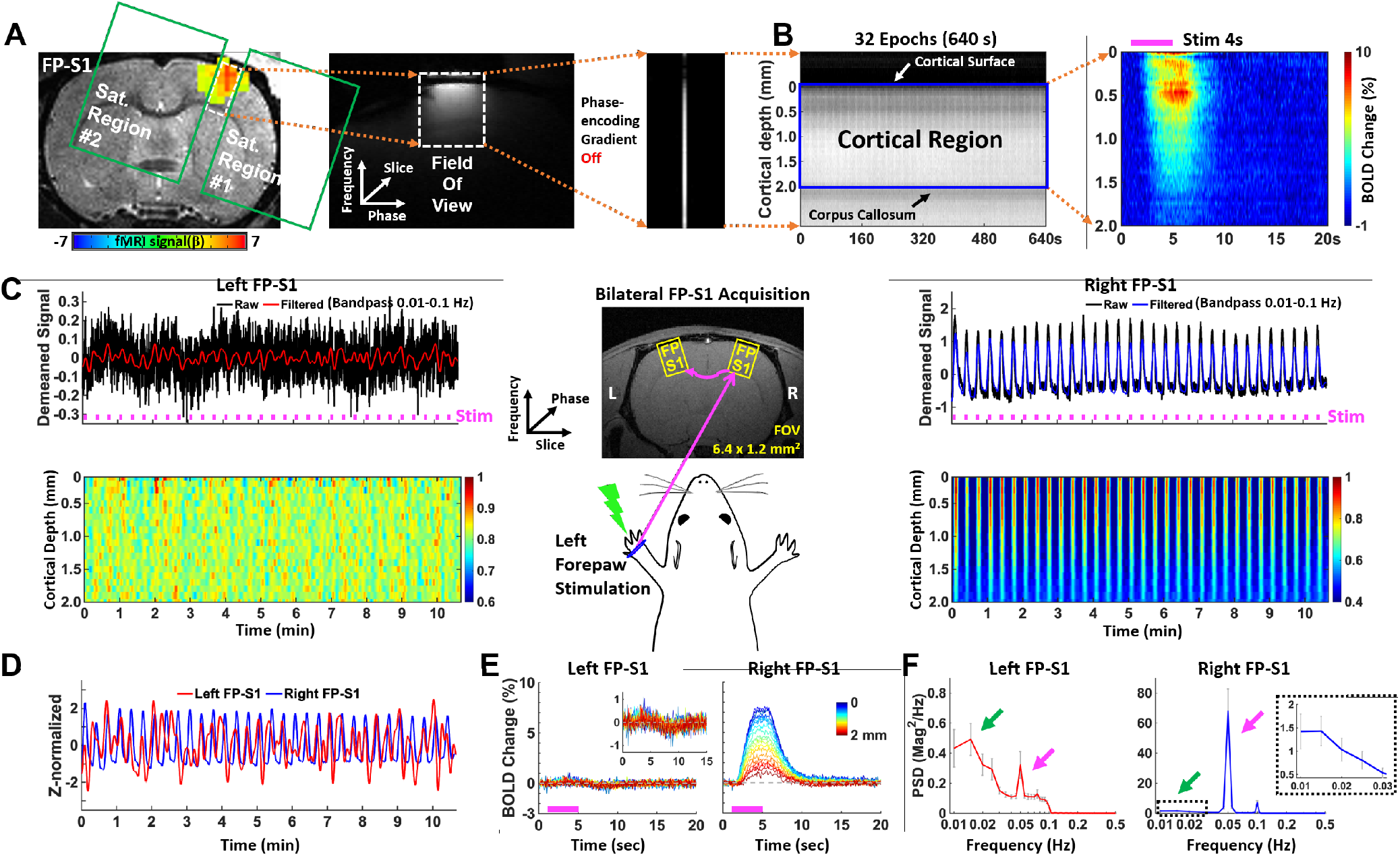
Evoked BOLD responses upon left forepaw stimulation using the unilateral and bilateral line-scanning method. **A-B.** Sequential procedure to acquire the average BOLD change map in the unilateral line-scanning acquisition. **A.** The procedure to set up the line-scanning method. *Left:* EPI BOLD activation map of forepaw somatosensory cortex overlaid on an anatomical RARE image. GLM-based t-statistics in AFNI is used, p <0.001. *Middle:* Representative reduced FOV (6.4 × 1.2 mm^2^) image with 2 saturation slices. *Right:* Single line-scanning profile acquired without phase-encoding gradient. **B.** *Left:* Spatiotemporal map concatenated with multiple line-scanning profiles for 32 epochs (10 min 40 sec). *Right*: Average BOLD percentage changes (block design: 1 s pre-stim, 4 s stim on and 15 s post-stim) show the laminar-specific BOLD responses across the cortical depth (0-2 mm, 50 μm resolution) from the cortical surface (white arrow) to the surface of the corpus callosum (black arrow). (**C-F**) Group-averaged results in bilateral line-scanning acquisition (n = 23 trials of 4 rats).**C.** *Top-left and-right:* Demeaned fMRI time series (32 epochs, 10 min 40 sec) of raw (black) and filtered (red and blue) data (average of 20 voxels, bandpass: 0.01-0.1 Hz) in the left and right FP-S1 regions during electrical stimulation (3 Hz, 4 s, 2.5 mA) to left forepaw. *Bottom-left and-right:* Normalized spatiotemporal maps show the laminar-specific responses along cortical depth in the left and right FP-S1 (0-2 mm, 100 μm resolution). *Middle*: Schematic illustration of the bilateral line-scanning experimental design in the coronal view of the symmetric FP-S1. **D.** The Z-score normalized fMRI time series (average of 20 voxels) of the left (red) and right (blue) FP-S1 show the stimulation-induced activation. **E.** Average BOLD time courses across the cortical depth (0-2 mm, 20 lines in total) show the evoked BOLD responses in the left and right FP-S1. **F.** The PSDs of the filtered fMRI time series (average of 20 voxels) of the left (red) and right (blue) FP-S1, showing the very low-frequency responses (0.01-0.02 Hz, green arrow, dashed black box for right FP-S1) and stimulation-induced peaks (0.05 Hz, magenta arrow). Error bars represent mean ± SEM across 4 animals.

### Mapping the bilateral laminar BOLD responses based on CC activation

Using BiLS method, we focused on analyzing the interaction of laminar-specific BOLD fMRI signals through callosal projections with the left forepaw stimulation. We created a 2D line profile map to cover both gray matter (cortex, FP-S1) and white matter (corpus callosum, CC). As shown in **Fig. 2A–B**, the grayscale functional maps demonstrated different T2*-weighted signal intensity of cortical and CC regions given their different T2* values, presenting the dynamic fMRI responses in the color maps (details in **Method**). From the representative trial, we extracted the Z-score normalized fMRI signals from FP-S1 and CC of two hemispheres, highlighting the salient evoked BOLD signal in the right FP-S1, as well as the ultra-slow oscillation patterns (0.01-0.02 Hz) in both sides (**Fig. 2C-F**). Moreover, PSD analysis revealed the power peak at the stimulation frequency of 0.05 Hz in both FP-S1 regions, showing much higher power for the right FP-S1 than the left FP-S1 (**Fig. 2G**). Interestingly, the 0.05 Hz peak also existed in bilateral CC regions, demonstrating evoked BOLD signals in the white matter of the representative trial (**Fig. 2H**). However, it was noteworthy that not all trials show robust 0.05 Hz peaks through transcallosal projections. **Fig. S2** showed another representative trial with strong evoked BOLD signals in the right FP-S but not detectable BOLD signals in the left side despite the similar ultra-slow oscillatory patterns across two hemispheres. This method allowed us to identify trial-specific interhemispheric laminar interactions driven by callosal projection neurons [60, 61, 67].

**Figure 2.**
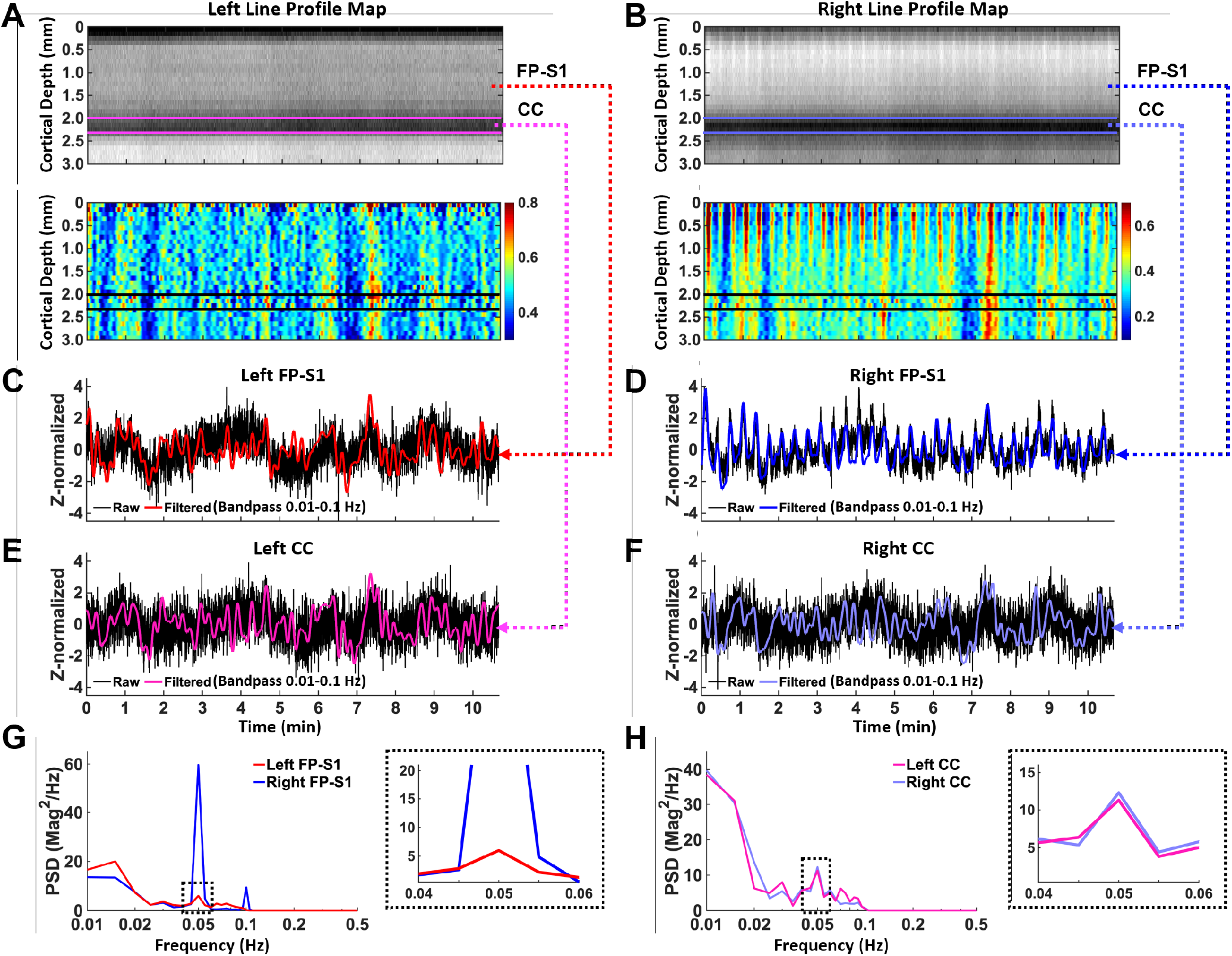
Evoked fMRI time series and BOLD responses in bilateral FP-S1 and corpus callosum regions from a representative trial. **A** and **B.** *Top:* Spatiotemporal maps consist of bilateral line-scanning profiles which were concatenated for 32 epochs (10 min 40 sec) from the left and right FP-S1 (0-2 mm), the left and right corpus callosum (2.0-2.3 mm) between the magenta lines and between the light purple lines respectively. *Bottom:* Normalized spatiotemporal maps show the laminar-specific responses in cortex and corpus callosum responses for the same regions as the upper images. The black lines indicate the left and right corpus callosum regions, the same as the magenta and light purple lines of the top images. **C** and **D.** The Z-score normalized fMRI time series of raw (black) and filtered (red and blue) data (average of 20 voxels, bandpass: 0.01-0.1 Hz) in the left and right FP-S1 during electrical stimulation to the left forepaw (block design: 1 s pre-stim, 4 s stim, and 15 s post-stim).**E** and **F.** The Z-score normalized fMRI time series of raw (black) and filtered (magenta and light purple) data (average of 20 voxels, bandpass: 0.01-0.1 Hz) in the left and right corpus callosum with the same period as **C** and **D**. **G** and **H.** The power spectral densities (PSDs) of the filtered and Z-score normalized fMRI time series from the left (red) and right (blue) FP-S1 (**G**), left (magenta) and right (light purple) corpus callosum regions (**H**) clearly show the evoked frequency responses (0.05 Hz) with enlarged PSDs (0.04-0.06 Hz, dashed black box).

Next, we investigated the CC activation-dependent laminar-specific interactions among different experimental trials. Based on the BOLD signals detected in the CC ipsilateral to stimulation, we sorted all the trials into two groups, *i.e.,* with or without the 0.05Hz peak in the PSD. As shown in **Fig. 3A**, Group 1 (10 trials) showed peaked power at 0.05 Hz with normalized values at 0.59 ± 0.17 (mean ± STD), which was significantly higher than other trials in Group 2 (13 trials, 0.26 ± 0.06) without a detectable peak. **Fig. 3B** showed the color-coded power spectral maps across different cortical layers, demonstrating a salient peak at 0.05 Hz in the right FP-S1 of both groups (**Fig. 3B**, white arrows, **Fig. S3 B and D**); however, the 0.05 Hz peak was only detected in the left FP-S1 of Group 1, but not in Group 2 (**Fig. 3B**, black arrows). In contrast, the ultra-slow oscillation at 0.01-0.02 Hz in both FP-S1 regions were reliably detected in both groups (**Fig. 3B**), implying a CC-independent global neuromodulation effect.

**Figure 3.**
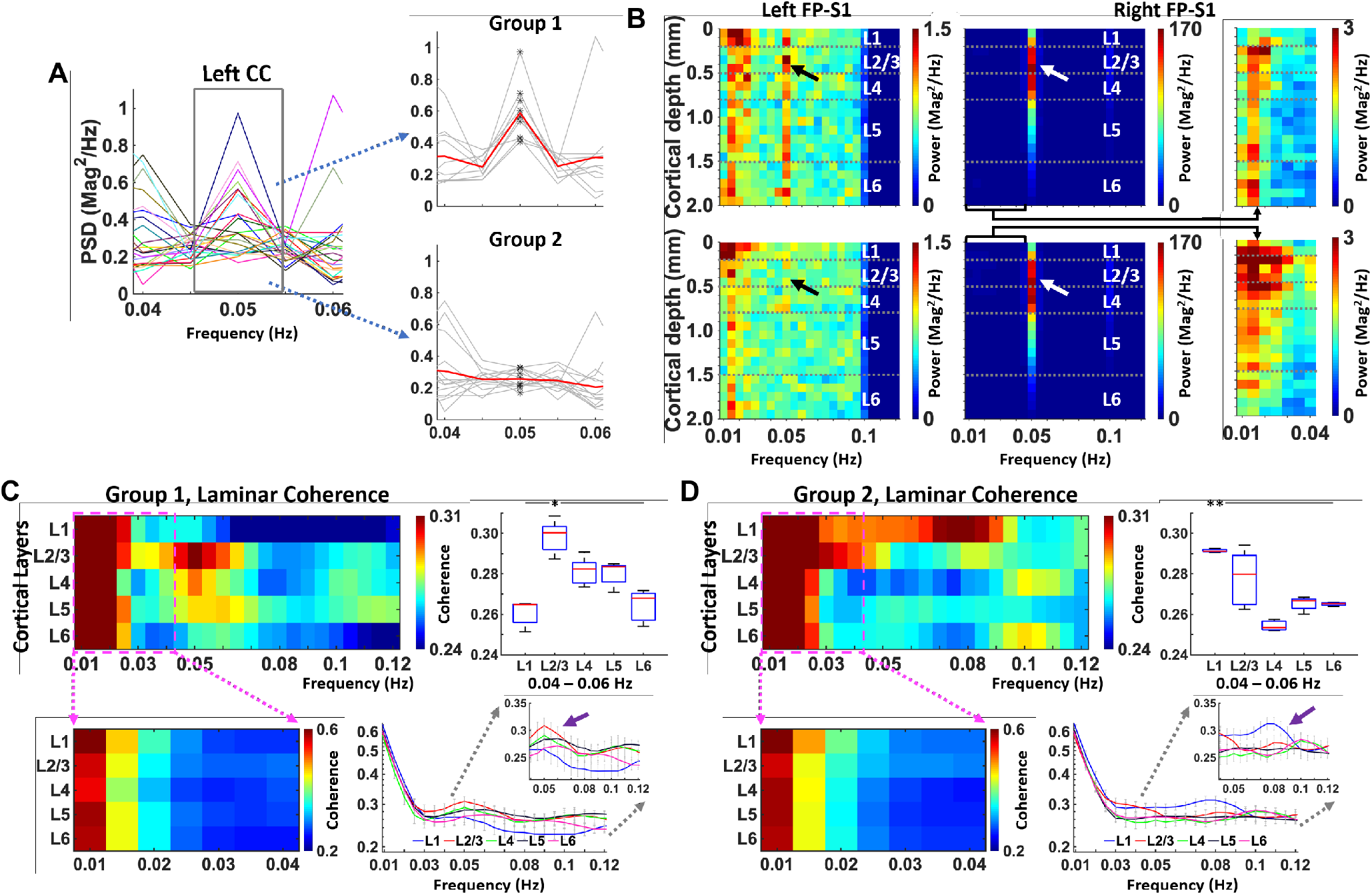
Corpus callosum activation-based grouping to characterize laminar-specific coherence patterns. **A.** *Left:* The PSD plots (0.04-0.06 Hz) of the filtered fMRI time series (average of 3 voxels, bandpass: 0.01-0.1 Hz) in the left corpus callosum to identify the stimulation frequency (0.05 Hz) (n = 23 trials of 4 rats). *Right:* The corpus callosum activation-based grouping showing Group 1 (upper) and Group 2 (lower) with and without the significantly high stimulation frequency (Student t-test: p=1.4156*10^-6), respectively (red line: mean, gray lines: individual trials, black points: values at 0.05 Hz).**B.** *Left and middle:* The voxel-wise PSDs of the filtered time series (20 voxels, bandpass: 0.01-0.1 Hz) in the left and right FP-S1 from the Group 1 (upper, 10 trials) and Group 2 (lower, 13 trials), respectively. *Right:* The enlarged voxel-wise PSDs (0.01-0.04 Hz) in the right FP-S1 region of the corresponding groups. **C** and **D.** *Top-left:* Group-averaged results representing laminar-specific coherence across the layers (L1, L2/3, L4, L5, L6) in Group 1 (**C**) and Group 2 (**D**). *Top-right:* the quantitative comparison of the laminar-specific coherences at 0.04-0.06 Hz. L2/3 is significantly different from the other layers (one-way ANOVA: p = 2.9262e-07, post-hoc: *p <0.05, Bonferroni correction) in©oup 1 (**C**) meanwhile L1 is significantly different from the other layers (one-way ANOVA: p = 1.6276e-07, post-hoc: **p <0.05, Bonferroni correction) in group 2 (**D**). *Bottom-left:* the enlarged view of coherence in the low-frequency at 0.01-0.04 Hz to clearly show the laminar-specific coherence patterns of Group 1 (**C**) and Group 2 (**D**). *Bottom-right:* Average layer-wise coherences© Group 1 (**C**) and group 2 (**D**). Error bars represent mean ± SEM across 4 animals. The purple arrows indicate the layer-dependent coherence values at the stimulation frequency (0.05 Hz).

Furthermore, we performed laminar-specific Z-score normalized PSD analysis, showing that L2/3 had a strong peak at 0.05 Hz in the left FP-S1 of Group 1, but not in Group 2 (**Fig. S3 A and C**). Meanwhile, laminar-specific coherence analysis between two FP-S1 regions revealed higher coherence coefficients at the 0.01-0.02 Hz were across all the cortical layers in both groups, but the coherence coefficients peaked at 0.05 Hz in Group 1 were primarily located at L2/3 which was significantly higher than other layers (**Fig. 3C**). As previously reported, 80% of callosal projection neurons (CPN) are located in L2/3 [61], explaining the strong coherence in the L2/3 of Group 1. Also, noteworthy was that the coherence coefficients of Group 2 showed higher values in L1 than other layers, but not specific to 0.05 Hz (**Fig. 3D**), which might be caused by the fMRI signal fluctuation dominated by larger draining veins at the cortical surface. In summary, these results not only demonstrated the coexistence of the ultra-slow oscillation and layer-specific bilateral BOLD correlation, but also highlighted the CC-dependent laminar-specific interhemispheric activation patterns upon stimulation.

### Mapping laminar-specific bilateral functional connectivity at different frequencies during rest

Besides CC activation-dependent interhemispheric interaction, we investigated the laminar-specific bilateral rs-fMRI signals in anesthetized rats. **Fig. 4A** showed representative Z-score normalized time courses from bilateral FP-S1, as well as 2D line profile rs-fMRI maps. The bilateral fMRI signal fluctuation (0.01-0.1 Hz) was highly correlated, and the oscillatory signals of both FP-S1 were peaked at 0.01-0.02 Hz and 0.08-0.1 Hz in PSD plots (**Fig. 4B**). Laminar-specific coherence analysis of all datasets revealed strong coherence at 0.01-0.02 Hz across all layers and significantly higher coherence coefficients of 0.08-0.1 Hz were detected at L2/3 than other layers (**Fig. 4C**). This result further confirmed the CC-based L2/3-specific rs-fMRI signal fluctuation at 0.08-0.1 Hz.

**Figure 4.**
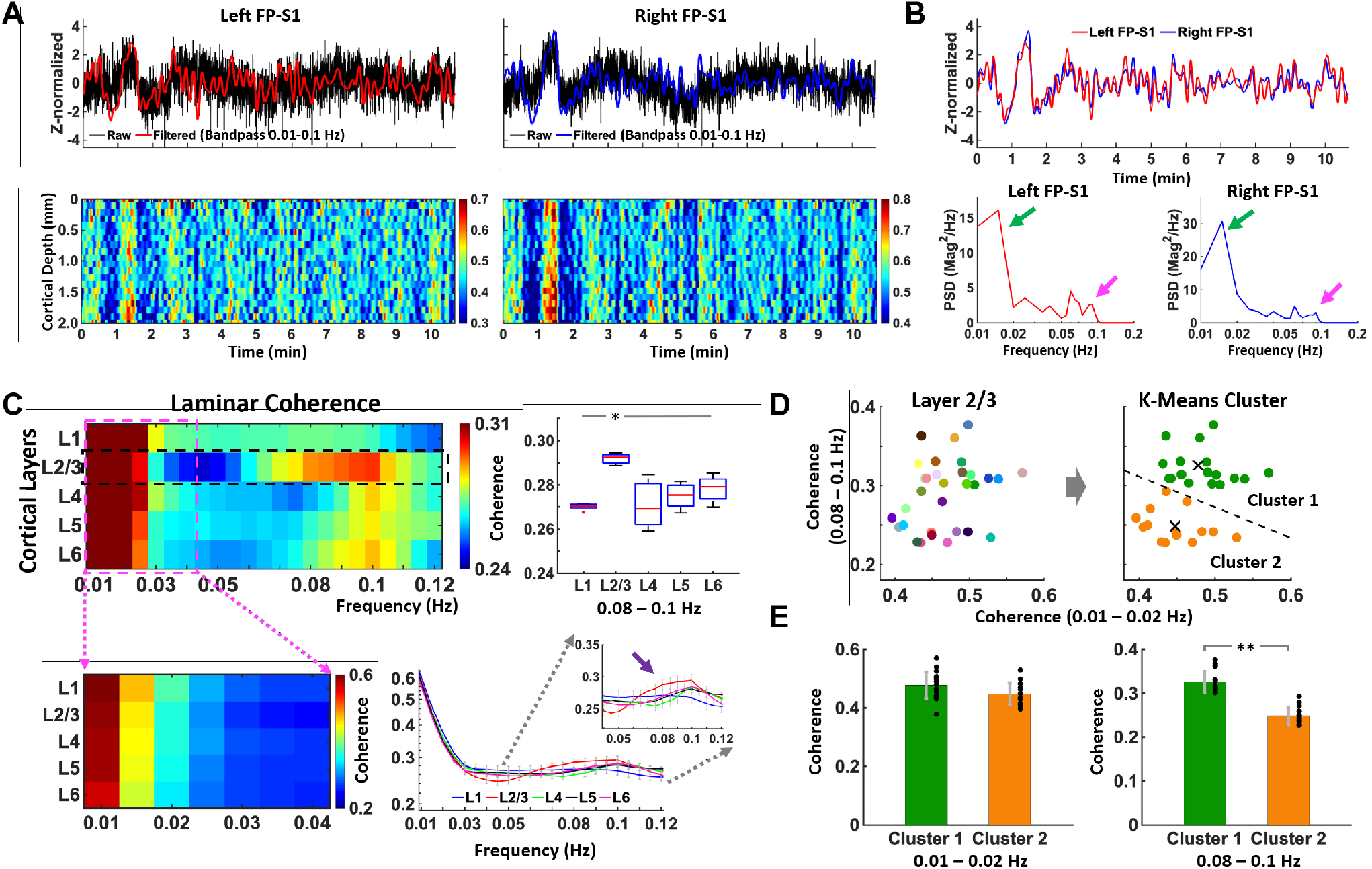
Characteristics of laminar-specific coherences in rs-fMRI using BiLS method. **A.** *Top:* Z-score normalized fMRI time series of raw (black) and filtered (red and blue) data (average of 20 voxels, 0-2 mm, bandpass: 0.01-0.1 Hz) in the left and right FP-S1 during rest (10 min 40 sec, one representative trial). *Bottom:* Normalized spatiotemporal maps showing the laminar-specific responses across the cortical depth in the same FP-S1 as the upper images (0-2 mm, bandpass: 0.01-0.1 Hz). **B.** *Top:* Z-score normalized fMRI time series of the filtered data from the same trial in the left and right FP-S1 marked in red and blue, respectively. (**A**). *Bottom:* The PSD analyses of the Z-score normalized time series in both the left and right cortices show obvious peaks at the ultra-slow oscillatory frequency (green arrow, 0.01-0.02 Hz) and 0.08-0.1 Hz (magenta arrow). **C-E.** Average results of coherence from all data sets (n = 32 trials of 4 rats) **C.** *Top-left:* Laminar-specific coherence across L1, L2/3, L4, L5, L6 (n = 32 trials of 4 rats). *Top-right:* the average coherences (0.08-0.1 Hz) across the whole trials showing that L2/3 was significantly different from the other layers (one-way ANOVA: p = 1.3477e-04, post-hoc: *p <0.05, Bonferroni correction). *Bottom-left:* the enlarged view of coherence at 0.01-0.04 Hz to clearly show the laminar-specific coherence patterns. *Bottom-right:* Average layer-wise coherences. Error bars represent mean ± SEM across 4 animals. The purple arrow indicates the layer-dependent coherences in the low-frequency (0.08-0.1 Hz). **D.** *Left:* Scatterplot of the L2/3 coherence values with x-axis as the mean of coherence at the very low-frequency (0.01-0.02 Hz) and y-axis as the mean of coherence at the low-frequency (0.08-0.1 Hz) to examine the dependence of the two different frequency bands. *Right:* By applying the PCA-based k-means clustering with the L2/3 coherence values, all the 32 trials were divided into two groups. Cluster 1 (n=18 trials) had relatively high averaged coherence values in both the frequency bands while Cluster 2 (n=14 trials) had low averaged coherence values. ‘×’ signs indicate the centroid of the individual groups. **E.** Cluster 1 and 2 show no significant difference at 0.01-0.02 Hz *(left,* independent t-test, p = 0.05464) while significant difference at 0.08-0.1 Hz *(right,* independent t-test, **p = 1.4127*10^-10).

To examine the relationship between L2/3-specific (0.08-0.1Hz) and ultra-slow rs-fMRI signal fluctuation, we plotted the trial-by-trial coherence coefficients at 0.01-0.02 Hz and 0.08-0.1 Hz (**Fig. 4D**). The large variability of coherence across all trials enabled the clustering analysis to specify the dependency of two frequency-specific oscillatory features. For example, **Fig. S4** showed another representative trial with lower power of ultra-slow oscillation but higher power of L2/3-specific fluctuation at 0.08-0.1 Hz. We applied PCA-based k-means clustering method (**Fig. 4D, S4 C and E**) to divide all trials into two clusters. Despite similar trial coherence distribution at 0.01-0.02 Hz, there were significantly different coherent coefficients of 0.08-0.1 Hz between the two clusters (**Fig. 4D, E** and **S4 D**). These results further demonstrated independent neuronal regulatory sources contributing to bilateral rs-fMRI signal fluctuations at 0.01-0.02 Hz across all the cortical layers, indicating a global neuromodulation effect, and at 0.08-0.1 Hz specific to L2/3, which can be modulated by the callosal projection neurons connecting two hemispheres.

## DISCUSSION

We applied the BiLS method to investigate circuit-specific interhemispheric interaction of bilateral fMRI signals in anesthetized rats. Laminar-specific coherence analysis demonstrates two independent low-frequency oscillatory patterns. One is the 0.01-0.02 Hz ultra-slow oscillation across all cortical layers in both stimulation and resting conditions, which may be caused by global neuromodulation. The other is the L2/3-specific coherence pattern, which appeared at the 0.05 Hz according to the stimulation paradigm with repetitive epochs (one epoch for 20 s duration), or 0.08-0.1 Hz during rest. The L2/3-specific coherence pattern in both conditions indicates that the laminar-specific interhemispheric interaction is likely associated with intrinsic neuronal activities mediated by callosal projection neurons.

Two challenging issues have limited the laminar-specific FC studies in rodent brains. First, fMRI is typically performed with a spatial resolution of several hundreds of microns [68–71], and the conventional EPI-based fMRI is often performed with the temporal resolutions of 1-2 s [72–74]. The limited spatial or temporal resolution makes it difficult to elucidate circuit-specific interhemispheric FC. Even though very high spatial resolution can be achieved at the cost of a slow sampling rate [34][75–77], the rs-fMRI signal is possibly confounded by the aliased cardiorespiratory artifacts [48, 63–66]. Secondly, the high temporal signal to noise ratio (tSNR) is needed for correlation/coherence analysis of rs-fMRI signal for FC mapping [78–80]. In contrast to task-related fMRI experiments with massive averaging to increase the tSNR, it remains a challenge to achieve sufficient tSNR for rs-fMRI with high spatiotemporal resolution at a single trial. Also, trial-by-trial variability will be pivotal to be verified when statistical analysis is used to extract the general traits of the connectivity pattern based on the neuronal source [81]. To overcome these problems, the line-scanning scheme provides a critical mapping strategy to record laminar fMRI signals along the cortical depth in high spatiotemporal resolution and with sufficient tSNR [82].

Technical advances at the ultra-high magnetic field *(e.g.*, 14 T) in combination with focal MRI signal acquisition provide an excellent opportunity to study the laminar-specific interhemispheric FC. As shown in **Fig. 2**, the gray matter (0-2 mm) and corpus callosum (2.0-2.3 mm) of FP-S1 were identified with the BiLS method. It is noteworthy that the temporal resolution of BiLS allows to eliminate the potential artifacts by temporal band-passed filters without concerning aliasing effect (**Fig. 2C)** [1, 83]. The artifact caused by cardiac pulsations are usually linked to large arteries located at the cortical surface [63, 64, 84]. As represented in **Fig. S1**, the fast-sampling line-scanning scheme enables to detect laminar-specific pulsation-related artifacts (*i.e.,* the 5-8 Hz cardiac cycles in rodents) [52], which are primarily located at L1. The lack of confounding artifacts to the laminar-specific rs-fMRI signal, especially at cortical layers deeper than L2/3, provides a unique platform to investigate the circuit-specific interhemispheric FC. Also, the laminar coherence map of Group 2 highlighted the superficial layer (L1) (**Fig. 3D**), which may involve the non-neuronal noises from cardiovascular responses. Meanwhile, given gradient-echo (GRE) based MR sequences, large draining veins located in superficial layer result in higher signal fluctuation and spreading temporal responses which may also contribute to this observation [36].

Despite extensive studies to elucidate the transcallosal-mediated interhemispheric neuronal interactions [8, 9, 60, 85, 86], CC-dependent interhemispheric rs-fMRI FC are primarily verified based on loss-of-functional studies in acallosal patients [9–12] or animals with callosotomy [13–15]. The observation of maintaining interhemispheric FC in several rs-fMRI studies [9–11] raised the need to specify circuit-specific rs-fMRI signal fluctuations, in particular, when originating from varied neuronal sources. Using the BiLS method, we decomposed the bilateral low frequency rs-fMRI signal fluctuation with laminar specificity, revealing two independent slow oscillatory patterns *(i.e.,* 0.01-0.02 Hz and 0.08-0.1 Hz) possibly driven by different regulatory mechanisms (**Fig. 4**). Callosal projection neurons are primarily located in L2/3 and project to both L2/3 and L5 of the opposite hemisphere [61, 87], which is well-matched with the detected laminar fMRI coherence peaks significantly located at L2/3, and partially at L5 (**Fig. 3C** and **4C**). Also, callosal circuit-specific optogenetic stimulation increased the power of the gamma frequency in L2/3 and L5 neurons [88], and mediated unique BOLD response patterns through ortho-and antidromic projections [89]. A future study to combine the callosal circuit-specific optogenetic stimulation with BiLS will further elucidate the causal relationship. In addition, robust ultra-slow fluctuation (0.01-0.02 Hz) across all the cortical layers with the BiLS method is consistent with the global BOLD signal fluctuation in anesthetized rats [90]. This global signal fluctuation has been correlated with the brainwide neuronal oscillatory sources in a brain state dependent manner across different species [90–93]. In particular, both subcortical neuronal projections through direct neuromodulation [22, 28, 94, 95] and autonomic regulation [96–98] could converge their regulatory effect to mediate the ultra-slow oscillatory patterns across cortical layers [25–30].

It should be noted that the nature of the local neuronal circuitry that generates interhemispheric fluctuations and modulates the frequency bands of rs-fMRI signals remains an open issue [6]. Nir and colleagues reported the low-frequency modulation in spontaneous firing rate and gamma LFP (40–100 Hz) as potential neural correlates of interhemispheric rs-fMRI signal fluctuation in the human sensory cortex [99]. Similarly, Mateo and colleagues also reported gamma-rhythm oscillations at ~0.1 Hz rhythm through vasomotion may serve as the underlying mechanism of the low-frequency hemodynamic signal fluctuation [6, 7]. This has been further specified with the arteriole specific single-vessel rs-fMRI study in anesthetized rats [62]. Moreover, hemodynamic signals correlate tightly with synchronized gamma oscillations in cats [100] and monkeys [101]. Interestingly, Schoelvinck and colleagues reported that the LFP power of upper gamma-range frequency (40-80 Hz) and lower frequency range at 2-15 Hz recorded from a single cortical site were positively correlated with global rs-fMRI signals [101]. This series of studies have provided strong evidence of the temporal neural correlate of rs-fMRI signal fluctuation. However, they do not assign the frequency-specific fluctuation to underlying neural projection circuits, which is critical to verify the distinct neural basis of FC. This study, for the first time, shows the frequency-specific oscillatory patterns across different cortical layers, presenting an important step to elucidate the neuronal circuit basis of the FC with rs-fMRI.

Several limitations about the BiLS method should be considered when interpreting the results of this study and for future optimization using high field rs-fMRI studies. Firstly, because of usage of saturation slices to delineate the FOV, adiabatic RF pulses are needed to produce sufficient and uniform signal saturation outside of the FOV. Alternative mapping strategy using 90-180° spin-echo-based line-scanning scheme will be further explored in multi slice line-scanning acquisition [82]. Secondly, due to the poor inhomogeneity of high magnetic field and different draining vein distribution on cortical surface across animals, tSNR can vary across line profiles from two hemispheres. To achieve sufficient tSNR, we can apply inductive coils [102] or wireless amplified NM detectors [103, 104] by region-specific implantation. Thirdly, although we only focused on the anesthetized rodent brains, the line-scanning method has been applied for human brain mapping [105, 106]. Despite cortical folds and fissures, it is possible to position the FOV to cover focal cortical regions to achieve laminar-specific fMRI signals with the multi-slice line-scanning scheme. Meanwhile, the multi-slice line-scanning scheme can be combined with the laminar electrophysiological recording or fiber photometry-based fluorescent recordings from genetically encoded biosensors, *e.g.* Ca^2+^, Glutamate, and dopamine [90, 107–109], to further elucidate the neuronal basis of the circuit specific rs-fMRI FC.

## METHODS

### Animal preparation

The study was performed in accordance with the German Animal Welfare Act (TierSchG) and Animal Welfare Laboratory Animal Ordinance (TierSchVersV). This is in full compliance with the guidelines of the EU Directive on the protection of animals used for scientific purposes (2010/63/EU). The study was reviewed by the ethics commission (§15 TierSchG) and approved by the state authority (Regierungspräsidium, Tübingen, Baden-Württemberg, Germany). A 12-12 hour on/off lighting cycle was maintained to assure undisturbed circadian rhythm. Food and water were available ad libitum. A total of 4 male Sprague-Dawley rats were used in this study.

Anesthesia was first induced in the animal with 5% isoflurane in the chamber. The anesthetized rat was intubated using a tracheal tube and a mechanical ventilator (SAR-830, CWE, USA) was used to ventilate animals throughout the whole experiment. Femoral arterial and venous catheterization was performed with polyethylene tubing for blood sampling, drug administration, and constant blood pressure measurements. After the surgery, isoflurane was switched off, and a bolus of the anesthetic alpha-chloralose (80 mg/kg) was infused intravenously. After the animal was transferred to the MRI scanner, a mixture of alpha-chloralose (26.5 mg/kg/h) and pancuronium (2 mg/kg/h) was constantly infused to maintain the anesthesia and reduce motion artifacts.

### EPI fMRI acquisition

All data sets from rats were acquired using a 14.1T/26 cm (Magnex, Oxford) horizontal bore magnet with an Avance III console (Bruker, Ettlingen) and a 12 cm diameter gradient system (100 G/cm, 150 μs rising time). A home-made transceiver surface coil with a 10 mm diameter was used on the rat brain in all experiments. For the functional map of BOLD activation (**Fig. 1A**), a 3D gradient-echo EPI sequence was acquired with the following parameters: TR/TE 1500/11.5 ms, FOV 1.92 × 1.92 × 1.92 cm^3^, matrix size 48 × 48 × 48, spatial resolution 0.4 × 0.4 × 0.4 mm^3^. A high order (*e.g.*, 2^nd^ or 3^rd^ order) shimming was applied to reduce the main magnetic field (B0) inhomogeneities at the region-of-interest. For anatomical reference of the activated BOLD map, a RARE sequence was applied to acquire 48 coronal images with the same geometry as that of the EPI images. The fMRI design paradigm for each trial comprised 200 dummy scans to reach steady-state, 10 pre-stimulation scans, 3 scans during stimulation, and 12 post-stimulation scans with a total of 8 epochs.

### BiLS acquisition

GRE-based BiLS datasets were acquired in anesthetized rats for evoked and rs-fMRI. BiLS was applied by increasing the slice dimension (1 to 2) to record fMRI signals in both left and right FP-S1 cortices while swapping the phase and slice encoding direction from the conventional line-scanning method, and by using two saturation slices to avoid aliasing artifacts in the reduced field-of-view along the phase encoding *(i. e.*, from rostral to caudal) direction (**Fig. 1A and C**; middle). The phase-encoding gradient was turned off to acquire line profiles (**Fig. 1A**, right). Laminar fMRI responses from the two cortices were acquired along the frequency-encoding direction (**Fig. 1C**, middle). The following acquisition parameters were used: TR/TE 100/12.5 ms, TA 10 min 40 sec, FA 45°, slice thickness 1.2 mm, slice gap 8.0 mm, FOV 6.4 × 3.2 mm^2^, and matrix 64 × 32. The fMRI design paradigm for each epoch consisted of 1 second pre-stimulation, 4 seconds stimulation, and 15 seconds post stimulation with a total of 20 seconds. A total of 6400 lines *(i.e.*, 10 m 40 s) in each cortex were acquired every single trial in evoked and rs-fMRI. Evoked BOLD activation was identified by performing electrical stimulation to the left forepaw (300 μs duration at 2.5 mA repeated at 3 Hz for 4 seconds). For anatomical reference of BiLS, the 2D gradient-echo (GRE) sequence was applied in each session to acquire 16 slices in the coronal plane with the following parameters: TR/TE 500/3.1 ms, FA 40°, slice thickness 0.5 mm, slice gap 0.5 mm, FOV 19.2 × 19.2 mm^2^, and matrix 192 × 192.

### Data Analysis

All signal processing and analyses were implemented in MATLAB software (Mathworks, Natick, MA) and Analysis of Functional NeuroImages software [110] (AFNI, NIH, USA). For evoked fMRI analysis for **Fig. 1A**, the hemodynamic response function (HRF) used was the default of the block function of the linear program 3dDeconvolve in AFNI. BLOCK (L, 1) computes a convolution of a square wave of duration L and makes a peak amplitude of block response = 1, with *g*(*t*) = *t*^4^*e*^-*t*^/[4^4^ *e*^-4^]. Each beta weight represents the peak height of the corresponding BLOCK curve for that class. The HRF model is defined as follows:

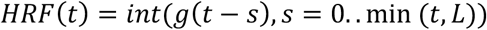

Cortical surfaces and CC were determined based on T2* contrast of fMRI line profiles (**Fig. 2A and B, Fig. S2 A and B**). The detailed processing was conducted as provided in the previous line-scanning study [38]. The line profile map concatenated with the multiple fMRI signals was normalized by a maximum intensity. The Z-score normalized time courses were calculated as follows: (x -μ)/σ, where x was original fMRI time courses and μ, σ were the mean and the standard deviation of the time courses, respectively (zscore function in MATLAB). Average BOLD time series and percentage changes were defined as (S-S0)/S0 × 100 %, where S was the BOLD signal and S0 was the baseline, *i. e.*, the average fluctuation signal in 1 second pre-stimulation window in evoked fMRI and mean epoch with 20 seconds (32 epochs for 640 s). The BOLD time series in each ROI were detrended and bandpass filtered (0.01-0.1 Hz, FIR filter) before analyzing PSD and coherence.

For PSD analysis, fMRI time series were used or converted to different forms depending on the purpose of power spectrum analysis. Original fMRI time series were used in **Fig. 1F** and **3B** to show different power amplitudes of evoked fMRI signals between both FP-S1 regions. In the representative trials (**Fig. 2** and **S2**), the FP-S1 and CC time series were converted to the Z-score normalized time series. The converted time series were used for PSD analysis to compare the frequency responses between the different regions avoiding the dependency of signal amplitudes between them (**Fig. 2G, H** and **Fig. S2 G, H**). For **Fig. 3A**, the demeaned fMRI time series of the individual trials were used. The PSD values from all the trials were down-or up-scaled to have the same mean at the two adjacent frequencies (*i.e.*, 0.045 and 0.055 Hz) of the stimulation frequency (0.05 Hz). Then, all the trials were divided into the two groups with the boundary lines (*i.e.*, mean of the PSD values at 0.05 Hz). PSDs were calculated by Welch’s power spectral density estimate method (pwelch function in MATLAB, FFT length: 2000, overlap: 50%).

Coherence analysis, which is generally accepted as an indicator of functional corticocortical connections, *e.g.*, increased coherence, is thought to reflect increased FC [111–113]. In this study, coherence analysis was employed as an essential indicator of laminar-specific interhemispheric FC and of which layer patterns between the symmetric cortices appeared in different ranges of low-frequency (<0.1 Hz). Layer-wise fMRI line profiles were used and converted to the Z-score normalized line profiles to minimize the dependency of the differences of baseline signal intensity between the left and right cortices. The PSDs of the Z-score normalized line profiles were computed for evoked fMRI as shown in **Fig. S3 A-D**. The coherence is defined as follows:

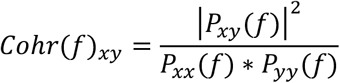

where *x,y* indicates the Z-score normalized fMRI time series from each cortical region, and *P_xx_*(*f*), *P_yy_*(*f*) is the individual PSD, and *P_xy_*(*f*) is the cross PSD. The coherence was calculated by ‘mscohere’ function in MATLAB (Hamming window length: 2560, FFT length: 2000, overlap: 50%). The frequency resolution of the coherence was 1/200 s = 0.005 Hz which provides the capability to observe slow oscillatory frequencies.

For clustering in rs-fMRI, the k-mean clustering method [114] was applied after preprocessing the coherence scatter points (**Fig. 4D**) via the principle component analysis (pca function in MATLAB) as shown in **Fig. S4 C** (kmeans function in MATLAB, MaxIter: 10000, Replicates: 10). The k-mean clustering converged under 10 iterations without tunning parameters.

For statistical analysis, one-way ANOVA was performed to compare the layer-wise coherence value using the post-hoc test in the evoked and rs-fMRI data. The coherence plot with error bars is displayed as the individual means ± SEM. An independent samples t-test was applied to compare the means of two independent groups and to determine whether there was a statistically significant difference between the associated population means. Student t-test was performed to calculate a p-value between group 1 and 2. The p-values <0.05 were considered statistically significant.

## Data availability

Excel files are included for each quantitative plot included in the main figures. All other data generated during this study are available from the corresponding author upon reasonable request.

## Code availability

The related image processing codes are available from the corresponding author upon reasonable request.

## ACKNOWLEDGEMENTS

This research was supported by NIH Brain Initiative funding (RF1NS113278-01, R01 MH111438-01), and the S10 instrument grant (S10 MH124733) to Martinos Center, German Research Foundation (DFG) Yu215/2-1, 3-1, BMBF 01GQ1702, and the internal funding from Max Planck Society. This project has received funding from the European Union Framework Programme for Research and Innovation Horizon 2020 (2014-2020) under the Marie Skłodowska-Curie Grant Agreement No.896245. We thank Dr. R. Pohmann, Dr. J. Engelmann, Dr. N. Avdievitch, and Ms. H. Schulz for technical support, Dr. P. Douay, Ms. R. König, and Ms. M. Pitscheider for animal support, the AFNI team for the software support.

---

**Figure S1.**
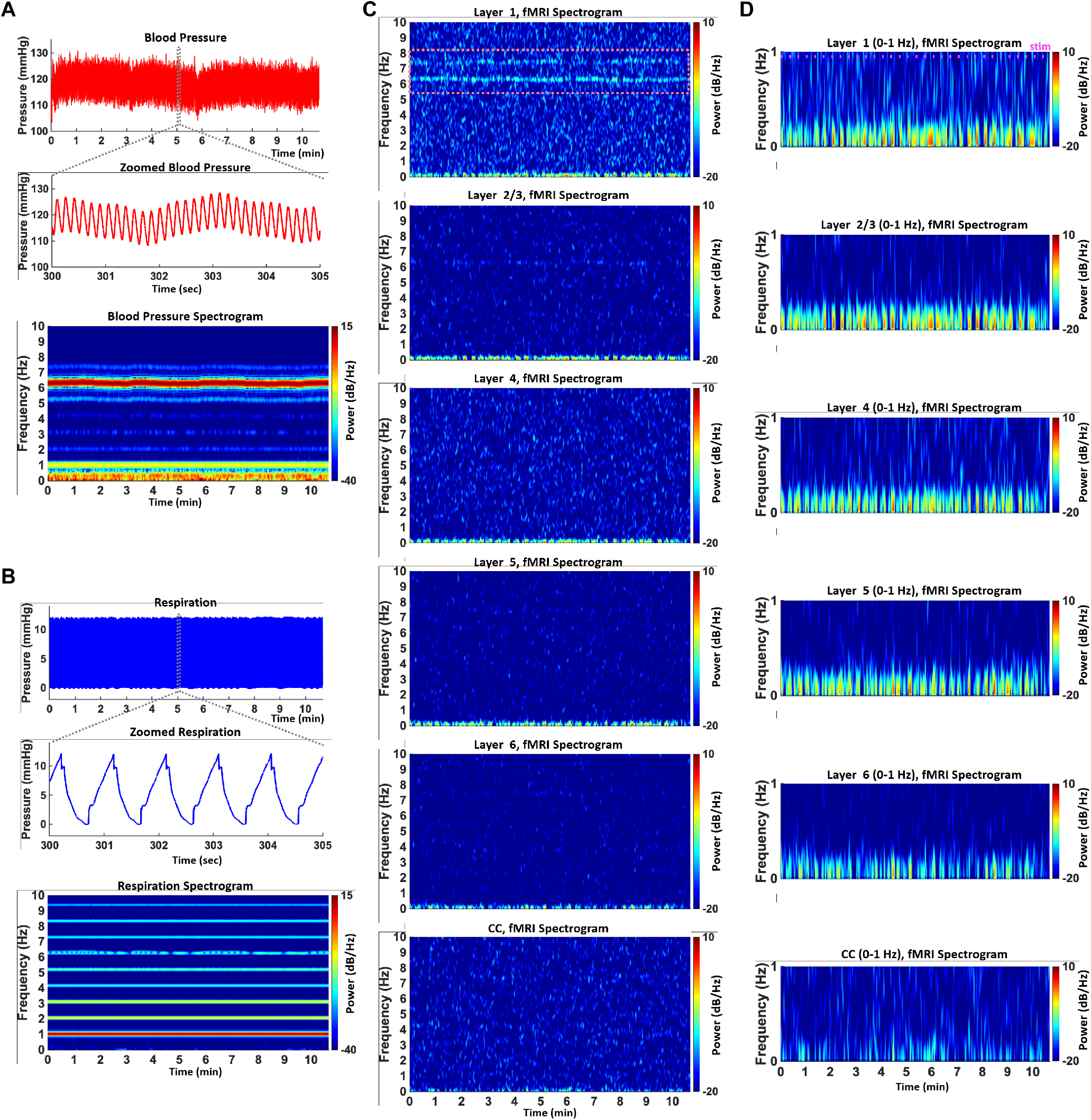
Effect of cardiorespiratory noises on laminar-specific evoked fMRI signals from a representative trial (TR 50 ms).**A**. *Top:* Arterial blood pressure time courses. *Middle:* Zoomed time courses from the upper time course (gray box). *Bottom:* The blood pressure spectrogram showing 6-7 Hz cardiac cycle. **B**. *Top:* Respiration time courses. *Middle:* Zoomed time courses from the upper time course (gray box). *Bottom:* The respiration spectrogram showing 1 Hz respiratory cycle and its harmonics. **B**. Layer-wise fMRI spectrograms from L1 to CC (from top to bottom) to show aliasing artifacts of the cardiorespiratory noises in the individual layers. **C**. Zoomed layer-wise fMRI spectrograms which show that no cardiorespiratory aliasing effects exist on evoked fMRI responses (0-1 Hz).

**Figure S2.**
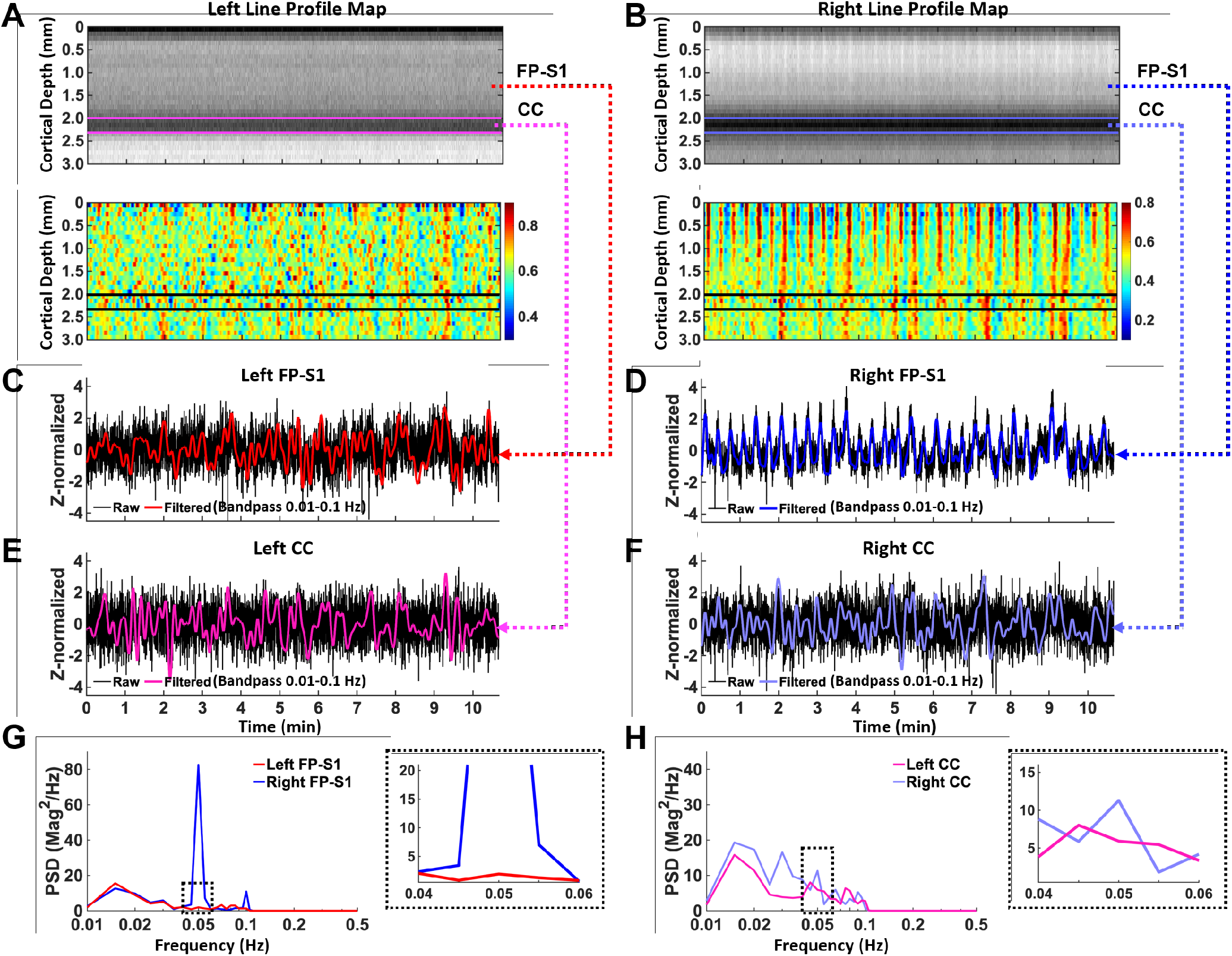
Evoked fMRI time series and BOLD responses in bilateral FP-S1 and corpus callosum from one representative trial (absence of CC activation).**A** and **B**. *Top:* Spatiotemporal maps consist of bilateral line-scanning profiles which were concatenated for 32 epochs (10 min 40 sec) from the left (red) and right (blue) FP-S1 (0-2 mm), the left and right corpus callosum (2.0-2.3 mm) between the magenta lines and between the light purple lines respectively. *Bottom*: Normalized spatiotemporal maps show the laminar-specific responses in the cortex and corpus callosum responses for the same regions as the upper images. The black lines indicate the left and right corpus callosum regions, the same as the magenta and light purple lines of the top images. **C** and **D**. The Z-score normalized fMRI time series of raw (black) and filtered (red and blue) data (average of 20 voxels, bandpass: 0.01-0.1 Hz) in the left and right FP-S1 regions during electrical stimulation to the left forepaw (block design: 1 s pre-stim, 4 s stim, and 15 s post-stim, 32 epochs, 10 min 40 sec).**E** and **F**. The Z-score normalized fMRI time series of raw (black) and filtered (magenta and light purple) data (average of 20 voxels, bandpass: 0.01-0.1 Hz) in the left and right corpus callosum regions during the same period as **C** and **D**. **G** and **H**. The power spectral densities (PSDs) of the filtered and Z-score normalized fMRI time series show the evoked frequency responses (0.05 Hz) with enlarged PSDs (0.04-0.06 Hz, dashed black box) from the left (red) and right (blue) FP-S1 (**G**). No local peak exists at evoked frequency responses (0.05 Hz) in the left corpus callosum (**H**, magenta line).

**Figure S3.**
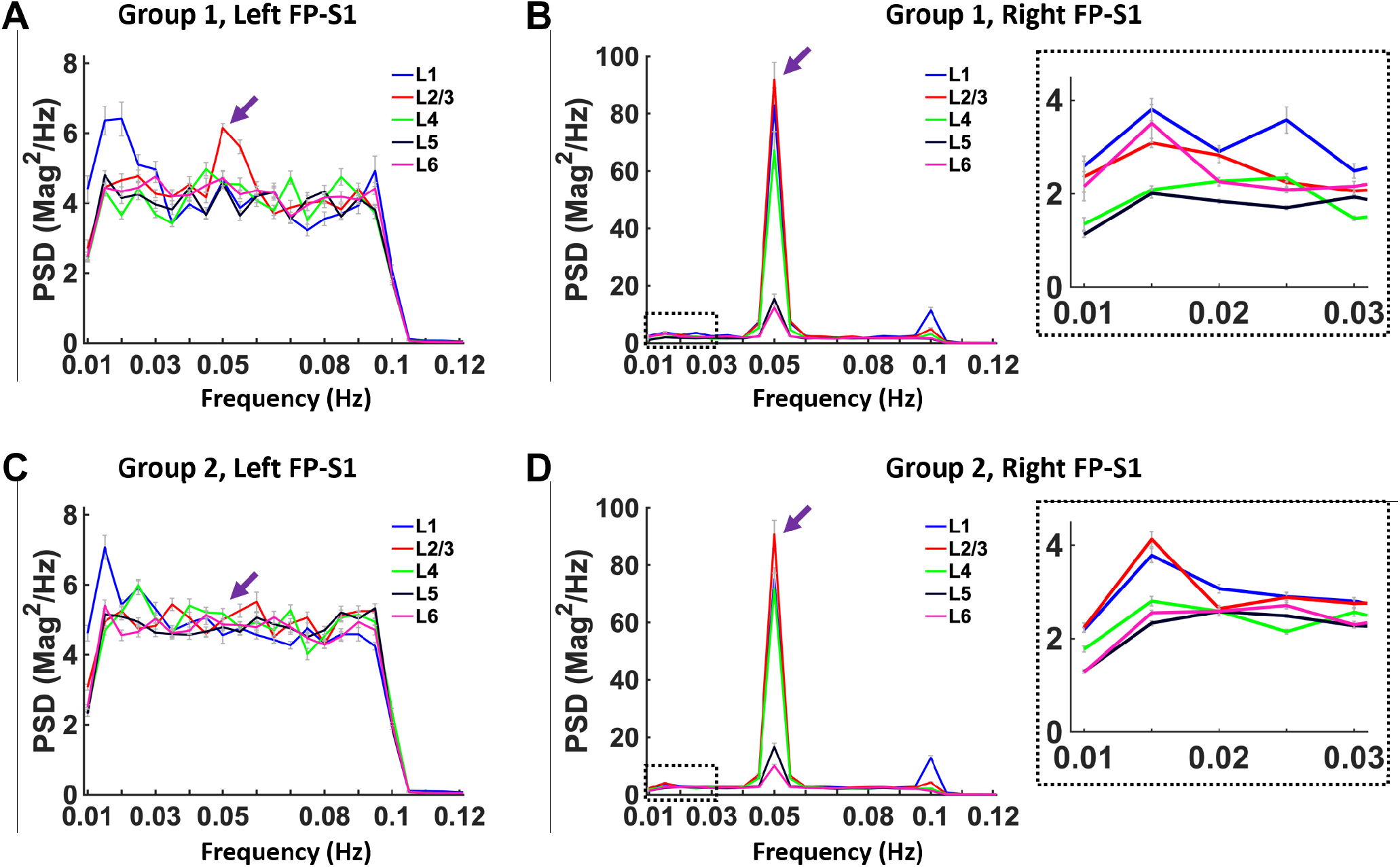
Layer-specific PSDs of the filtered and Z-score normalized line profiles as the input signals for coherence calculation in Group 1 and 2. **A** and **B**. Layer-specific PSDs of the filtered and Z-score normalized line profiles (Layer (L) 1-6, bandpass: 0.01-0.1 Hz) in the left (**A**) and right (**B**) FP-S1 regions of Group 1 (10 trials) with enlarged PSDs (0.01-0.03 Hz, dashed black box in **B**). The purple arrows indicate local peaks at the stimulation frequency (0.05 Hz) in both the left and right FP-S1 and the Layer 2/3 in the left FP-S1 has the highest PSD value (**A**, red line).**C** and **D**. Layer-specific PSDs of the filtered and Z-score normalized line profiles (L1-6, bandpass: 0.01-0.1 Hz) in the left (**C**) and right (**D**) FP-S1 regions of Group 2 (13 trials) with enlarged PSDs (0.01-0.03 Hz, black box in **D**). The purple arrows indicate local peaks at the stimulation responses (0.05 Hz) in the right FP-S1 regions, but not in the left side (**C**).

**Figure S4.**
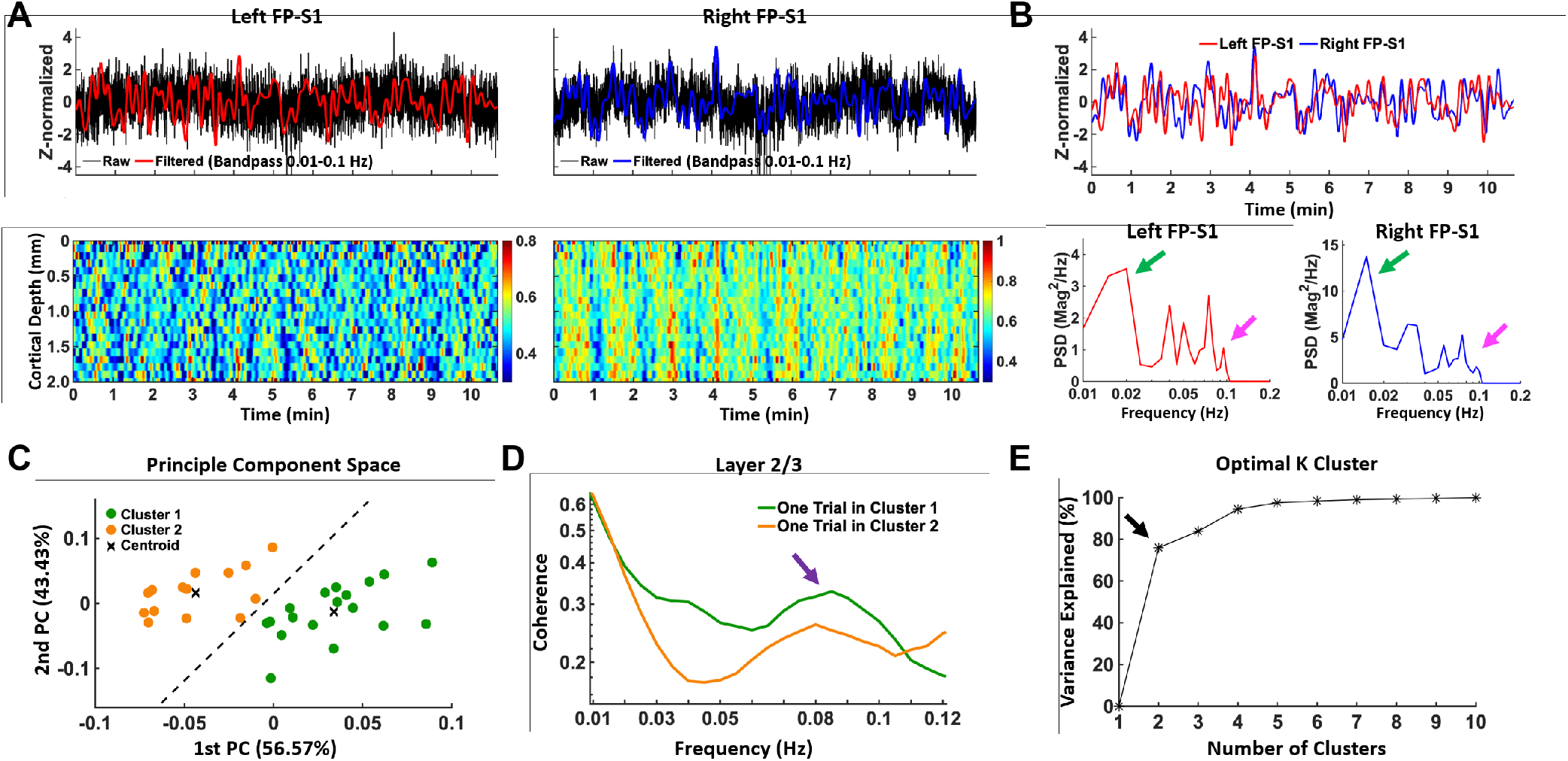
Resting-state (rs-) fMRI responses in bilateral FP-S1 regions from one representative trial (less synchronization) 704 and PCA-based k-means clustering in rs-fMRI. **A.** *Top:* Z-score normalized fMRI time series of raw (black) and filtered (red and blue) data (average of 20 voxels, 0-2 mm, bandpass: 0.01-0.1 Hz) in the left and right FP-S1 regions during rest (10 min 40 sec, one single trial). *Bottom:* Filtered (0.01-0.1 Hz) and normalized spatiotemporal maps show the laminar-specific responses across the same FP-S1 regions as the upper images. **B.** *Top:* The Z-score normalized fMRI time series of the filtered data (average of 20 voxels, 0-2 mm, bandpass: 0.01-0.1 Hz) in both the left and right FP-S1 regions (red and blue) from **A.** *Bottom:* The PSDs of the Z-score normalized time series show peaks at the ultra-slow fluctuation frequency (green arrow: 0.01-0.02 Hz) and 0.08-0.1 Hz (magenta arrow) in left (red) and right (blue) FP-S1. C. Scatterplot of L2/3 coherence values (Fig. 4D) by applying PCA-based k-means clustering. All the 32 trials were divided to two groups (Cluster 1, 2: n=18, n=14, respectively). ‘x’ signs indicate the centroid of the individual groups. D. L2/3 coherences with the single respective trial of Cluster 1 (green) and 2 (orange). The purple arrow indicates one trial in Cluster 1 had higher coherence values at 0.08-0.1 Hz than one trial in Cluster 2. E. Plot of the optimal number of clusters with the percentage of variance explained. The black arrow indicates number 2 was chosen as the optimal number of clusters in the elbow curve.

